# Natural Secretory Immunoglobulins Enhance Norovirus Infection

**DOI:** 10.1101/253286

**Authors:** Holly Turula, Juliana Bragazzi-Cunha, Sadeesh Ramakrishnan, Carol Wilke, Mariam Gonzalez-Hernandez, Alexandra Pry, Julianne Fava, Sophia Svoboda, Yatrik Shah, Blaise Corthesy, Bethany Moore, Christiane E. Wobus

## Abstract

Secretory immunoglobulins (SIg) are a first line of mucosal defense by the host. They are secreted into the gut lumen via the polymeric immunoglobulin receptor (pIgR) where they bind to antigen and are transported back across the FAE via M cells. Noroviruses are highly prevalent, enteric pathogens that cause significant morbidity, mortality and economic losses worldwide. Murine norovirus (MNV) exploits microfold (M) cells to cross the lymphoid follicle-associated epithelium (FAE) and infect the underlying population of immune cells. However, whether natural, innate SIg can protect against norovirus infection remains unknown. To investigate the role of natural SIg during murine norovirus pathogenesis, we used pIgR-deficient animals, which lack SIg in the intestinal lumen. Contrary to other enteric pathogens, acute MNV replication was significantly reduced in the gastrointestinal tract of pIgR-deficient animals compared to controls, despite increased numbers of dendritic cells, macrophages, and B cells in the Peyer’s patch, established MNV target cell types. Also, natural SIg did not alter MNV FAE binding or FAE crossing into the lymphoid follicle. Instead, further analysis revealed enhanced baseline levels of the antiviral molecules interferon gamma (IFNɣ) and inducible nitric oxide synthase (iNOS) in the small intestine of naive pIgR-deficient animals compared to controls. Removing the microbiota equalized IFNɣ and iNOS transcript levels as well as MNV viral loads in germ-free pIgR KO mice compared to germ-free controls. These data are consistent with a model whereby SIg sensing reduces pro-inflammatory, antiviral molecules, which facilitates intestinal homeostasis but thereby promotes MNV infection. In conclusion, these findings demonstrate that natural SIg are not protective during norovirus infection in mice and represent another example of indirect modulation of enteric virus pathogenesis by the microbiota.

## Introduction

The extensive mucosal surface of the gastrointestinal tract is a potential entry point for many pathogens. To protect itself from pathogen attack, the host has evolved multiple mechanisms, including the secretion of immunoglobulins (i.e., secretory Ig (SIg). SIg neutralize microbes in the intestinal lumen and groom enteric bacteria by mediating removal of pathogenic species and reducing the immunogenicity of remaining populations (Kaetzel 2014). Intestinal epithelial cells transcytose polymeric IgA (pIgA) and pIgM from the lamina propria via the basolaterally expressed polymeric immunoglobulin receptor (pIgR). Once the pIgR:pIgA/M complex reaches the intestinal lumen, the receptor is cleaved, and SIgA and SIgM are formed (Kaetzel 2014). This transcytotic process is used as a host defense to expel pathogens that have crossed the epithelial barrier and for pathogens present in intestinal epithelial cells (Corthesy 2009). Specifically, pIgR is necessary for the defense against such divergent enteric pathogens as Salmonella (Wijburg, Uren et al. 2006), Giardia muris (Davids, Palm et al. 2006), and rotavirus (Schwartz-Cornil, Benureau et al. 2002) and deletion of pIgR results in increased pathogen loads.

SIgA represent the predominant species of immunoglobulins in the intestine (Pabst, Cerovic et al. 2016) and as such are a critical first line of defense at mucosal surfaces (Corthesy 2007). However, in addition to their host defense function they also play an immunomodulatory role (Corthesy 2007). The follicle-associated epithelium (FAE) of Peyer’s patches (PP) and other mucosal-associated lymphoid follicles contain microfold (M) cells. M cells specialize in the uptake of luminal micro- and macromolecules across the FAE into PP to be sampled by the underlying antigen-presenting cells (APCs) (Corthesy 2007). SIgA aid in this luminal sampling and the initiation of mucosal immune responses. Selective adherence of luminal SIgA to the M cell surface triggers uptake of SIgA-immune complexes into the PP (Mantis, Cheung et al. 2002), resulting in ‘retrograde’ SIgA sampling by dendritic cells (DCs) (Rey, Garin et al. 2004). This uptake initiates non-inflammatory activation of DCs and induction of regulatory T cells (Corthesy 2007). The SIgA-induced anti-pathogenic immune responses also non-specifically reduce the inflammatory potential of macrophages via upregulation of inhibitory receptors (Mantis, Rol et al. 2011).

Noroviruses are the leading cause of acute gastroenteritis worldwide with an annual estimated global cost of $60 billion (Bartsch, Lopman et al. 2016). Every year, these viruses are responsible for the death of ~200,000 children under the age of 5 in developing countries and an estimated 21 million infections in the USA alone (Patel, Widdowson et al. 2008, Hall, Lopman et al. 2013). Yet, no approved virus-specific prevention and treatment strategies exist for this emerging pathogen. Noroviruses replicate in the gastrointestinal tract and infect multiple different cell types of both intestinal epithelial and immune cell origin *in vitro* and *in vivo* (Karst and Wobus 2015, (Karandikar, Crawford et al. 2016, Grau, Roth et al. 2017). Thus, these viruses must navigate the complex mixture of microbial and host-derived substances in the intestinal lumen to cross the intestinal epithelial barrier and establish an infection. Targeting these very early events during infection may provide an avenue for intervention, as they are instrumental in determining host range, initiation of immune responses, and pathogenesis. However, very limited information is available about factors that promote or inhibit the early steps during norovirus pathogenesis.

To gain a better understanding of the early events in norovirus infection, we and others use murine norovirus (MNV), an established and highly tractable animal model for studying norovirus biology (Karst and Wobus 2015). MNV is a natural mouse pathogen endemic to animal facilities across the world (Hsu, Wobus et al. 2005, Muller, Klemm et al. 2007, Kitajima, Oka et al. 2009, Kim, Lee et al. 2010). The first MNV strain, MNV-1, was discovered in 2003, but multiple MNV strains have been isolated since (Karst, Wobus et al. 2003, Thackray, Wobus et al. 2007). MNV-1 causes an acute infection and initiates infection in the ileum (Gonzalez-Hernandez, Liu et al. 2014), the distal most portion of the small intestine, which also has the greatest density of PP (Mowat and Agace 2014). It is in intestinal lymphoid follicles, such as PP, that MNV infection is primarily detected in antigen-presenting cells (APCs) (DCs, macrophages) and lymphocytes (T cells, B cells) (Wobus, Karst et al. 2004, Jones, Watanabe et al. 2014, Grau, Roth et al. 2017). To reach these target cells underneath the intestinal epithelium, MNV hijacks M cells and uses these cells as an entry portal for targeted delivery of virions to permissive cells (Gonzalez-Hernandez, Liu et al. 2013, Gonzalez-Hernandez, Liu et al. 2014, Karst and Wobus 2015). In addition to M cells, enteric bacteria are also required for optimal MNV pathogenesis (Jones, Grau et al. 2015, Baldridge, Turula et al. 2016). Despite the multi-faceted role of SIg in grooming luminal bacteria, defense from pathogens, and sampling of these complexes by M cells, nothing is known about the role of natural SIg during norovirus infection.

Thus, the goal of this study was to investigate the role of natural SIg in MNV-1 infection. Towards that end, we used pIgR-deficient animals (pIgR KO), which lack SIg in the intestinal lumen. Surprisingly, and contrary to the known protective role of natural SIg (Wijburg, Uren et al. 2006), MNV-1 viral loads were reduced in the gastrointestinal tract of pIgR KO animals compared to C57BL/6 controls (WT B6). This was despite enhanced numbers of DCs, macrophages, and B cells in the PP of pIgR KO mice. Inhibition of virus infection may have been in part due to enhanced baseline levels of interferon gamma (IFN-ɣ), a cytokine known to inhibit MNV translation (Changotra, Jia et al. 2009), and the antiviral molecule inducible nitric oxide synthase (iNOS), in naive pIgR-deficient animals compared to WT animals. Removing the microbiota adjusted IFN-ɣ and iNOS levels in germ-free pIgR KO to similar levels as in germ-free WT B6 animals and restored viral loads to WT levels in pIgR KO mice. Taken together, these data demonstrate that natural SIg do not play a protective but a proviral role during norovirus infection. These findings further suggest a model whereby SIg sensing reduces levels of some antiviral molecules and contributes to intestinal homeostasis, which in turn promotes MNV infection.

## Materials and Methods

### Animals

C57BL/6 animals were purchased from Jackson Laboratories (Bar Harbor, ME). A breeder pair of PIgR knockout mice on a C57BL/6 background (B6.129P2-*Pigr^tm1Fejo^*/Mmmh, stock number: 030988-MU) was obtained from Drs. Stappenbeck and Virgin (Washington University, St. Louis, MO). All animals were bred and housed in a SPF-free and MNV-free room. Serum tested negative for MNV using our ELISA(Wobus, Karst et al. 2004). All animals were used between 6 and 12 weeks of age. Animal studies were done in accordance with local and federal guidelines. The protocol was approved by the University of Michigan Committee on Use and Care of Animals (UCUCA number PRO00006658).

### MNV Virus Stocks

The plaque-purified MNV-1.CW3 (GV/MNV1/2002/USA) virus stock (herein referred to as MNV-1) was generated as previously described (Taube, Perry et al. 2012). A neutral red-labeled MNV-1 stock (MNV-1-NR) (3.8x10^6^ PFU/mL) was generated from MNV-1, passage 6, as previously described (Gonzalez-Hernandez, Perry et al. 2013). MNV-1-NR was handled in a darkened room using a red photolight (Premier OMNI, #SL-23R). 1 ml aliquots were stored in a light safe box at -80°C.

### Animal Infection

Animals were infected in a darkened room using a red photolight via oral gavage (o.g.) with 3.8x10^5^ plaque-forming units (PFU) of MNV-1-NR in 100 μl/mouse. Tissues were harvested at 9 or 18 hours post infection (hpi) as previously described (Gonzalez-Hernandez, Liu et al. 2014) with the following modifications: 2 cm of tissue was collected in pre-weighed 2 ml O-ring tubes (Denville Scientific #C19042-S) containing 1.0 mm diameter zirconia/silica beads (BioSpec #11079110z), flash frozen in an ethanol/ dry ice bath, weighed, and stored at -80°C.

### Plaque Assay

Homogenized samples were exposed to white light for 30 min to inactivate input virus. Light exposure reduced MNV-1-NR titers by 3 logs. The plaque assay was performed as previously described (Gonzalez-Hernandez, Bragazzi Cunha et al. 2012). Samples without detectable replicated virus were assigned 10 PFU/mL as the lowest detectable unit. Data was normalized to PFU per gram of tissue. The limit of detection (LOD) was determined for each organ by averaging the PFU/gram tissue of all samples without detectable viral titers.

### RNA Isolation

Total RNA was extracted from tissue samples via TRIzol Reagent (ThermoFisher Scientific) following manufacturers guidelines and resuspended in 30 μl RNase, DNase free water (Corning). Contaminating DNA was removed by treating samples with Turbo DNA free DNase kit (ThermoFisher Scientific). One μl TURBO DNase was added for 30 min at 37°C, before adding another 1 μl and incubating for an additional 30 min at 37°C. DNAse activity was stopped by adding 0.2 volumes of DNAse Inactivation Reagent for 5 minutes at room temperature. Total RNA was quantified using a spectrophotometer (NanoDrop #ND-1000) and stored at -80°C for further use.

### Quantitative Real-Time PCR

To measure host cell transcripts, cDNA was generated with 100 μg of total RNA using iScript Reverse Transcription Supermix for RT-qPCR (Bio-Rad) using the following protocol: 25°C at 5 min, 42°C at 30 min, 85°C at 5 min in a thermocycler (Eppendorf Mastercycler Epgradient PCR machine) and stored at -20°C. One μl of total cDNA was analyzed for levels of Gapdh (Wood, Rios et al. 2016), GP2 (Wood, Rios et al. 2016), Spi-B (Wood, Rios et al. 2016), IFN-ɣ (Mohanty, Shivakumar et al. 2006), interferon-beta (IFN-β) (Mohanty, Shivakumar et al. 2006), tumor necrosis factor-alpha (TNF-α) (Mohanty, Shivakumar et al. 2006), iNOS (Lopusna, Benkoczka et al. 2016Stoolman, Vannella et al. 2011), Interferon stimulated gene 15 (ISG15) (Lopusna, Benkoczka et al. 2016), transforming growth factor-beta (TGF-β) (Henderson, Mackinnon et al. 2006), interleukin-10 (IL-10) (Terashima, Watarai et al. 2008), interferon-lambda (IFN-λ) (Lopusna, Benkoczka et al. 2016) in a Biorad CFX96 Real Time System qPCR machine using Sso Advanced Universal SYBR Green Supermix (Bio-Rad). Gene expression was normalized to transcript levels of the endogenous host gene *Gapdh*, and fold change was calculated relative to WT C57BL/6 controls using the delta delta CT method as described (Higgins, Johnson et al. 2011). Quantification of MNV-1 genome equivalents in host tissue was performed as previously described (Taube, Perry et al. 2012).

### Ligated Intestinal Loops

Animals were anesthetized using 5% isoflourane and placed on a 37°C heating pad to maintain body temperature throughout the procedure to maintain body temperature throughout the procedure. The ileum was tied gently on either side of each PP using silk surgical suture thread (LOOK #SP118). Approximately 0.1 ml MNV-1 (6.7×10^7^ PFU/mL) was injected as the loose end of the loop was tightened, resulting in a 2 cm closed loop. The anesthetized mouse was monitored for 25 min, after which the PP were excised. Mucus was removed using 5 mM Dithiothreitol (Sigma-Aldrich) in 1x phosphate buffered saline (PBS) (Gibco) for 10 min in a 37°C water bath. The FAE was removed by shaking the PP at 250 rpm (MaxQ 600, Thermo Scientific) for 20 min in 1x PBS containing 5 mM EDTA (Fluka Analytical). PP were removed and free cells were collected. PP lamina propria and FAE were then washed twice with 1x PBS by gently inverting the tubes. Cells were placed in 500 μl RNALater (Sigma) and then stored at -80°C. For SIgA:MNV complex formation, MNV-1 (2.8×10^7^ PFU) was pre-incubated with 0.25 mg/ml recombinant SIgA (Crottet, Peitsch et al. 1999) (a generous gift from Dr. Blaise Corthésy, University of Lausanne) or 0.25 mg/ml bovine serum albumin fraction V (Roche #03116956001) diluted in 1x PBS for 1 hour at 37°C before adding to the loop and treated as described above.

### Tissue Digestion/ Flow Cytometry

PP and small intestine lamina propria were dissected from naive PIgR KO and WT B6 animals and digested as previously described (Geem, Medina-Contreras et al. 2012). Viable cells were enumerated using a hemocytometer and 0.4% trypan blue (Gibco). 2x10^6^ cells were then used for flow cytometric analysis. Cells were stained for cell viability (Invitrogen Live/Dead fixable Aqua Dead Cell stain Kit #L34957) for 15 min on ice in 1x PBS. Cells were then washed and stained for expression of the following surface markers: CD45 (diluted 1:200, clone 30-F11, AF700, Biolegend #103128), I-A/I-E (diluted 1:50, clone M5/114.15.2, PE/Dazzle 594, Biolegend #107647), CD19 (diluted 1:50, clone 6D5, PerCP/Cy5.5, Biolegend #115534), CD11b (diluted 1:50, clone M1/70 APC/Cy7, Biolegend #101226), CD64 (diluted 1:50, clone X54-5/7.1 FITC, Biolegend #139316) CD11c (diluted 1:50, clone N418 PE/Cy7, Biolegend #117318). All were diluted in FACS buffer; 5% fetal bovine serum (Hyclone #SH30396.03) and 0.1% sodium azide (Sigma) in 1x PBS, for 30 min on ice. After surface staining, cells were fixed for 10 min on ice using Cytofix/Cytoperm (BD Biosciences). After fixation, cells were washed and fluorescence was analyzed with a BD LSRfortessa (BD Biosciences). Data was analyzed using FlowJo software (BD Biosciences). The gating strategy was as follows. Live, CD45 and I-A/I-E (MHCII) positive APCs were identified and further characterized into B cells (CD19+), and mononuclear phagocytes (CD19-/CD11b+) were further delineated into macrophages via CD64 expression (CD64+/CD11c-), and DCs using CD11c (CD64-/CD11c+).

### Purification of MNV-1 capsid protein protruding (P) and shell (S) domains

P and S domain were bacterially expressed and recombinant proteins purified as previously described (Taube, Rubin et al. 2010).

### ELISAs

The SIgA ELISA was performed as described previously (Karst, Wobus et al. 2003) with the following modifications. Microtiter plates (Immulon II HB flat-bottomed ELISA plate, Thermo Labsystems) were coated with 0.1 mg/ml mouse SIgA, IgA dimer, or secretory component (kindly provided by Dr. B. Corthesy, Lausanne University) in coating buffer overnight at 4**°**C. Plates were washed and blocked overnight at 4**°**C. Recombinant P and S domain were then added (0.1 mg/ml) for 1 hour at 37**°**C. Bound MNV-1 capsid domains were detected with a rabbit polyclonal anti-MNV-1 VLP (diluted 1:5000) (Wobus, Karst et al. 2004) followed by a peroxidase-conjugated goat anti-rabbit secondary antibody (IgG diluted 1:1000, Jackson Laboratories) and absorbance was read at 405 nm. Mouse IgA was analyzed in feces and serum using the mouse-specific IgA kit following manufacturer’s protocol (Bethyl Laboratories, Montgomery, TX). For fecal IgA ELISA, 10% fecal slurry was made using the provided sample dilution buffer. The fecal slurry was further diluted 1:200 using sample dilution buffer and the IgA was assessed in 100 μl of the diluted slurry. For serum IgA analysis, 100 μl of 1:500 diluted serum samples (in sample dilution buffer) was used.

### Statistical Analysis

All data were analyzed using unpaired Student’s T test using the data analysis program, GraphPad Prism version 7 (GraphPad Software, La Jolla, CA). *= P<0.05. **= P<0.01 ***= P<0.001. ****= P<0.0001

## Results

### Acute norovirus infection is reduced in pIgR KO mice

To investigate the role of SIg during norovirus infection, we analyzed MNV-1 infection in mice lacking pIgR (Uren, Johansen et al. 2003), the receptor necessary for SIg transcytosis into the lumen. Since assessment of viral replication in the intestine is confounded by the lingering presence of the initial inoculum, neutral red, a light-sensitive dye, was incorporated into MNV-1 virions (MNV-1-NR) as previously described (Gonzalez-Hernandez, Perry et al. 2013). Exposure to light inactivates MNV-1-NR virions but not newly replicated MNV-1 particles devoid of NR, providing a means to differentiate between input (i.e., light-sensitive) and replicated (i.e., light-tolerant) virus (Gonzalez-Hernandez, Perry et al. 2013).

To determine the kinetics of acute MNV-1-NR infection, we first performed a time-course of MNV-1-NR infection in C57BL/6 mice. Mice were infected by oral gavage with 3.8x10_5_ PFU, and viral loads measured after 9, 12, 18, and 24 hpi. No replication was detected in the stomach and duodenum throughout the 24 hour time course (**Figure S1B–C**). Replicated viral titers were first detected at 9 hpi in the jejunum and ileum of C57BL/6 animals (**Figure S1D–E**). Ileal replicated viral titers peaked at 12 hpi and then plateaued, whereas jejunal replicated viral titers did not significantly increase until 24 hpi. A small amount of replicated virus was also detected at 9 hpi in the mesenteric lymph node (MLN) and increased steadily throughout the time course (**Figure S1A**). Replicated viral titers in the cecum and ascending colon were first detected 12 hpi and increased until 24 hpi (**Figure S1F–G**). Shedding of replicated virus was only observed in one animal at 18 hpi (**Figure S1H**). Taken together, these results suggest that the first round of MNV-1 replication occurs within 9 hpi in the distal small intestine after oral infection of C57BL/6 animals.

Next, we compared MNV-1 infection in naïve pIgR-deficient (pIgR KO) and C57BL/6 wild type (WT B6) mice. We first confirmed that fecal levels of IgA are undetectable in pIgR KO mice, while serum IgA levels are significantly increased compared to WT B6 animals (**Figure S2A-B**) in accordance with the literature (Johansen, Pekna et al. 1999, Shimada, Kawaguchi-Miyashita et al. 1999). Then, we measured viral loads in the intestinal tract and draining lymph nodes of pIgR-deficient (pIgR KO) and C57BL/6 wild type (WT B6) mice at 9 and 18 hpi, or after approximately one and two replicative cycles, respectively. Replicated viral titers were detectable at 9 hpi in the jejunum and ileum of pIgR KO and WT B6 animals (**Figure S3C–D**). Surprisingly, viral loads in the ileum of pIgR KO animals were significantly reduced compared to WT B6 controls (**Figure S3D**). No consistent infection was detected in the remaining organs in either group (**Figures S3A-B, S3E–G**). After a second round of MNV-1 replication at 18 hpi, however, replication was detected in most organs and the difference between pIgR KO and WT mice had magnified (Figure 1). Specifically, pIgR KO animals had significantly reduced levels of replicated viral titers in the MLN, jejunum, ileum, and colon compared to controls (Figure 1A, 1C–D, 1F). Replication was only detectable in a few animals in the duodenum in both groups (Figure 1B), while fecal shedding was only detectable in WT B6 mice (Figure 1G). The lack of fecal shedding in pIgR KO mice may be a reflection of the overall reduced virus load in these animals. Taken together, these data demonstrate that pIgR KO mice have reduced viral loads overall, suggesting that natural, non-MNV specific, SIg are not protective, but instead enhance acute norovirus infection *in vivo.*

**Figure 1:**
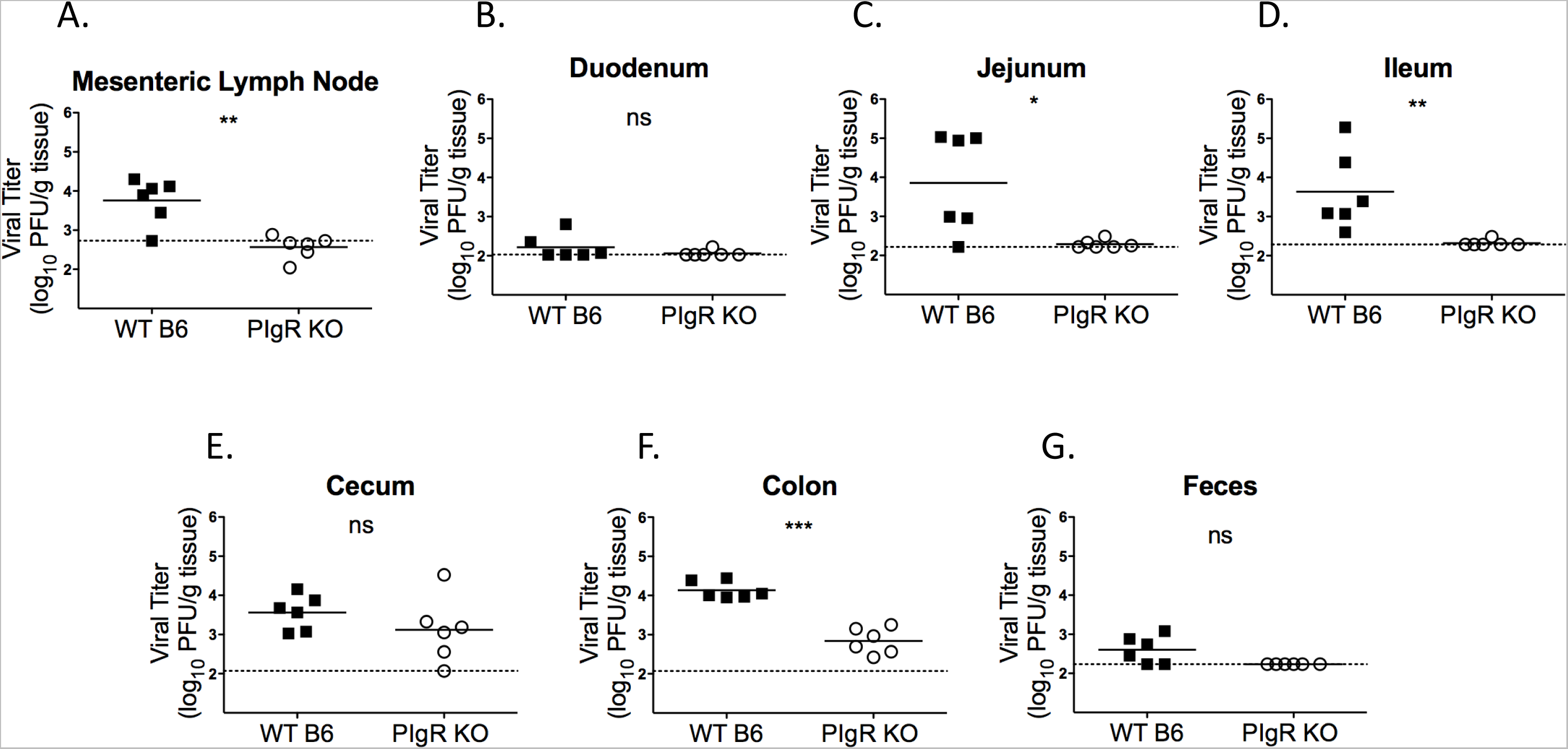
Acute norovirus infection is reduced in pIgR KO mice. **A-G.** Animals were each infected by o.g. with 3.8×10^5^ PFU/animal MNV-1-NR in 100 μl, and tissues were harvested at 18 hpi in a darkened room using a red photolight. The tissue homogenate was serially diluted and exposed to light for 30 min. Replicated viral titers in the indicated tissue of WT B6 and pIgR KO animals were assessed via plaque assay. The limit of detection for each graph was as follows (log PFU/g tissue): **A.** 2.728 **B.** 2.029 **C.** 2.219 **D.** 2.286 **E.** 2.071 **F.** 2.071 **G.** 2.235. Data are from two independent experiments and each symbol is a datapoint from an individual animal. Error bars are standard error of the mean (SEM). Data were analyzed using unpaired Student’s T test. *= P<0.05. **= P<0.01. ns = not statistically significant

### Natural Secretory Immunoglobulins do not aid in MNV-1 access to Peyer’s patches

To determine the mechanism of reduced MNV-1-NR infection of pIgR KO animals, we further characterized these mice. First, we investigated M cell-associated gene expression since M cells are necessary for efficient MNV infection (Gonzalez-Hernandez, Liu et al. 2013, Gonzalez-Hernandez, Liu et al. 2014). Host RNA was isolated from PP in the ileum of naïve pIgR KO and WT B6 mice and analyzed for transcript levels of SpiB and GP2, a transcription factor required during M cell development and a marker of mature M cells, respectively (Mabbott, Donaldson et al. 2013). No differences in expression levels were observed for both genes between both groups (Figure 2A). These data suggest that the virus encounters similar levels of M cells during infection of both mouse strains, and that the lack of pIgR and associated lack of SIg:antigen complex sensing does not significantly affect M cell development.

**Figure 2:**
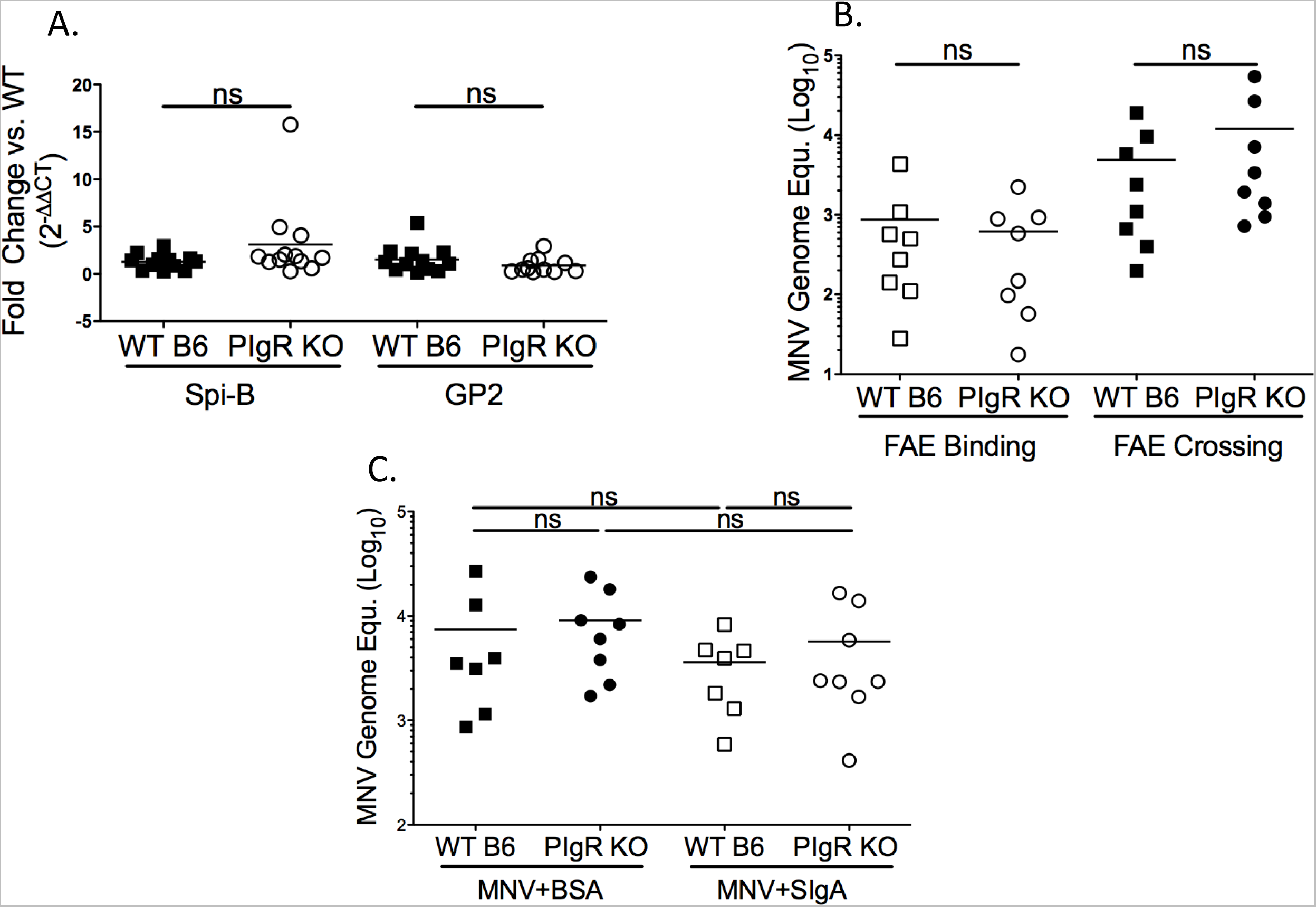
Natural secretory immunoglobulins do not aid in norovirus access to Peyer’s patches. **A.** Transcript levels of M cell-related genes, Spi-B and GP2, from WT B6 and pIgR KO Peyer’s patches over GAPDH transcript levels are shown as 2^∧^-(ΔΔCT) in reference to WT B6. ΔCT values were analyzed for statistical significance. **B-C.** Intestinal ileal loop assay in WT B6 and pIgR KO. 100 μl (3.81×10^5^ PFU) MNV-1 was injected into aclosed loop and incubated for 25 min. Viral genome copy equivalents were measured by RT-qPCR in Peyer’s patches. **B.** MNV-1 bound to the follicle-associated epithelium (FAE binding) and MNV-1 internalized into the Peyer’s patch (PP) lamina propria (FAE crossing). **C.** MNV-1 was pre-incubated with secretory IgA (SIgA) or bovine serum albumin (BSA) for 1 hour at 37°C before injecting the complex to the loop. MNV-1 genome equivalents per PP are shown. Data are from two to four independent experiments and each symbol is a datapoint from an individual PP. The data mean is indicated for each group. Data were analyzed using unpaired Student’s T test.ns = not statistically significant.

SIg are taken up by M cells and sampled by the underlying network of MNV-susceptible APCs (Corthesy 2007). To determine whether natural SIg aid in viral targeting to, and transcytosis across the epithelial barrier, we first determined whether MNV-1 is capable of interacting with non-NoV specific, recombinant sIgA. Recombinant anti-rotavirus SIgA, secretory component alone or dimeric IgA (Crottet, Cottet et al. 1999) were coated in ELISA plates and incubated with bacterially expressed MNV-1 P and S domains followed by detection with a polyclonal anti-MNV antibody (**Figure S4A**). The P domain of the MNV-1 capsid protein bound to sIgA as a whole and both components, the IgA dimer and secretory component. Binding of the S domain to SIgA or its components was similar to the negative control BSA. These data demonstrate the interaction with natural SIgA is restricted to the P domain, which also interacts with host cell receptors and contains neutralizing antibody epitopes (Kolawole, Smith et al. 2017). SIgA did not affect viral infection *in vitro* since no difference was observed in infectious MNV-1 titers by plaque assay in murine macrophages (RAW 264.7 cells) when complexed with recombinant, non-antigen specific SIgA compared to MNV-1 alone (data not shown).

Since MNV-1 could bind to recombinant SIgA, we next performed ligated ileal loop assays to determine whether MNV-1 uptake was altered by the presence or absence of SIg. Ileal PP of naïve pIgR KO and WT B6 animals were ligated to create a ~1 cm closed loop, and virus was injected as the loop was sealed. After 25 minutes, whole PP were excised and mucus-bound virions were removed via dithiothreitol (DTT) treatment prior to measuring viral genome titers by qRT-PCR. No difference was observed in viral genome copies (**Figure S4B**), suggesting similar amounts of virions bound and/or entered the PP in either mouse strain. However, MNV-1 infects primarily immune cells (Grau, Roth et al. 2017), so to investigate the amount of virions that reach the immune cells of the PP, we digested the FAE from the PP lamina propria with EDTA subsequent to DTT treatment and determined viral genome levels by qRT-PCR. Again, no differences were observed in the amount of viral genome associated with the PP FAE and those that crossed the FAE and associated with the lamina propria in pIgR KO versus WT B6 animals (Figure 2B). However, one limitation of the previous experimental design is the significantly reduced interaction time of MNV-1 with SIg compared to a natural, oral infection during which the virus travels through the intestinal tract. We hypothesized that pre-incubation of MNV-1 with natural SIgA may boost virus internalization if performed prior to the ileal loop experiments. However, we detected similar MNV genome equivalents in the PP lamina propria in WT B6 and pIgR KO animals when MNV-1 was pre-complexed with non-MNV specific SIgA or bovine serum albumin (BSA) as negative control (Figure 2C).

Taken together, these data demonstrate that although MNV-1 can interact with recombinant non-specific SIg, natural SIg does not facilitate MNV-1 binding to or crossing of the epithelial barrier. These findings indicate that natural SIg does not modulate norovirus interaction with the epithelial barrier.

### pIgR KO mice have increased numbers of small intestinal APC subsets

Since access to the site of viral replication, *i.e.,* PP, cannot account for the reduced infection seen in pIgR KO animals, we next investigated whether a reduced number or percent of B cells, macrophages and DCs, the known MNV target cell types at the time (Wobus, Karst et al. 2004, Jones, Watanabe et al. 2014), in the small intestine might provide an explanation. Towards that end, small intestines and their PP were collected from naïve animals. Tissues were digested to create single cell suspensions and cellularity was analyzed via flow cytometry. No difference was detected in the percentage of B cells (CD19+/CD11b-), macrophages (CD19-/CD11b+/CD64+), and DCs (CD19-/CD11b+/CD64-/CD11c+) in the villous or PP lamina propria (Figure 3A-B). No difference was observed in the number of the three APC subsets in the villous lamina propria (Figure 3C). Interestingly, all three APC subsets were increased roughly 2.5-fold in the PP lamina propria of pIgR KO animals compared to controls (Figure 3D). These data demonstrate that pIgR KO animals have increased numbers of MNV-susceptible cell types at the site of infection. Taken together with data from Figure 2, these data suggest that access to and availability of norovirus target cells cannot account for reduced MNV-1-NR infection seen in pIgR KO animals.

**Figure 3:**
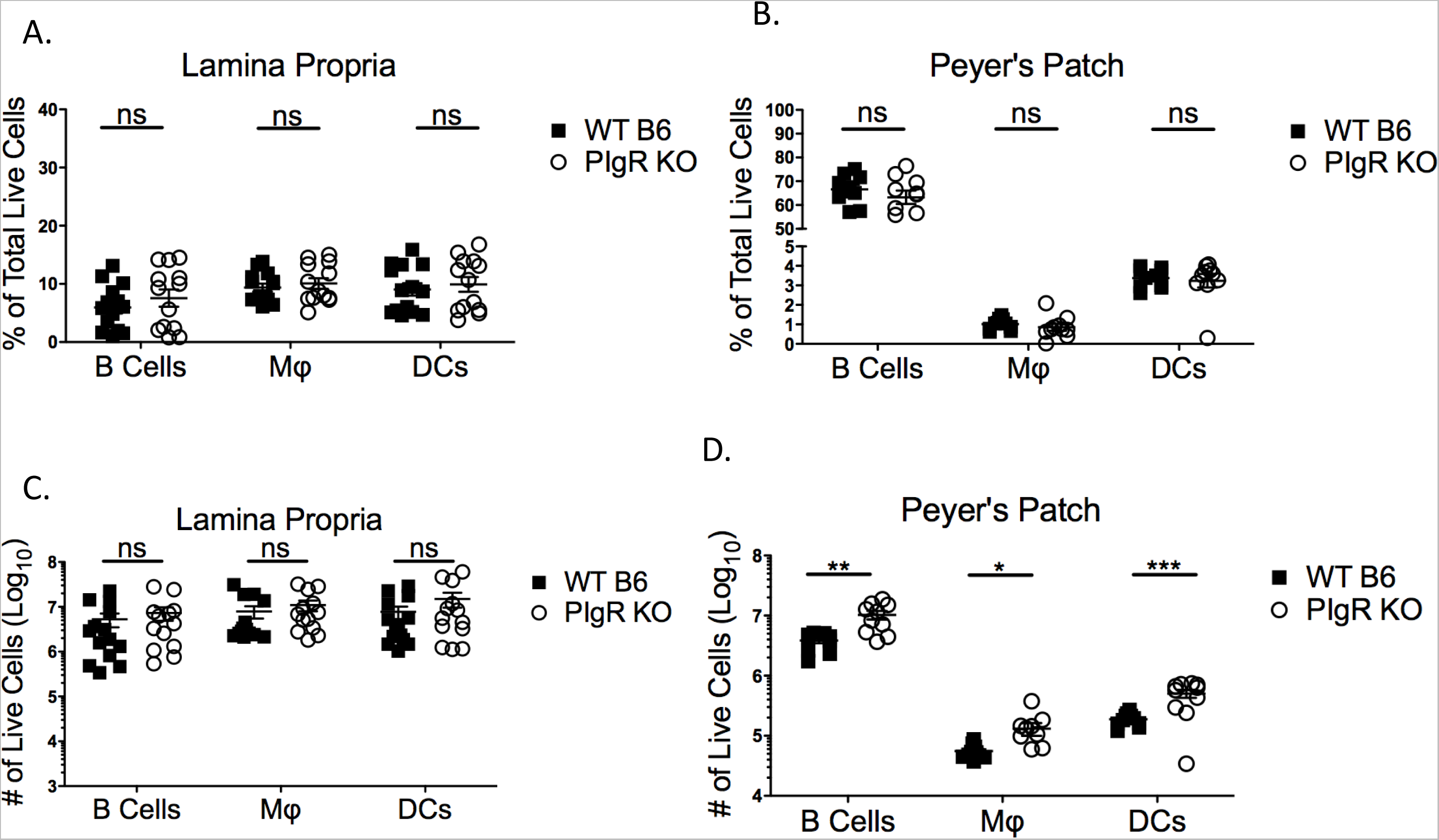
PIgR KO mice have increased small intestinal APC subsets. **A-D.** Peyer’s Patches and small intestinal lamina propriacells were isolated from naivepIgR KO and WT B6 animals. Live cells were analyzed via flow cytometry. The gating strategy was as follows: Antigen-presenting cells (APCs) were defined as Singlets/Live/CD45+/I-A/I-E (MHCII)+. APCs were further gated into B cells (CD19+/CD11b-), macrophages (CD19-/CD11b+/CD64+), and dendritic cells (CD19-/CD11b+/CD64-/CD11c+). **A-B.** Percentage of live lamina propria (**A**) and Peyer’s patch (**B**) cells. **C-D.** Absolute number of live lamina propria (**C**) and Peyer’s patch (**D**) cells. Data shown are from two to three independent experiments and each symbol is a datapoint from an individual animal. The data mean is indicated for each group. Data were analyzed using unpaired Student’s T test. *= P<0.05. **= P<0.01 ***= P<0.001. ****= P<0.0001. ns = not statistically significant.

### IFN-ɣ and iNOS levels are enhanced in naïve pIgR KO mice

SIg complexes have immunomodulatory functions, and when enteric bacteria are recognized in complex with SIg an anti-inflammatory response is initiated (Corthesy 2007). Therefore, we next investigated whether pIgR KO mice exhibit an altered intestinal immune landscape in the absence of SIg. Towards that end, transcript levels of a panel of cytokines and chemokines were analyzed in naïve pIgR KO and compared to those from WT B6 animals. The ileum was chosen for assessment, as it is the initial site of MNV-1 replication, and the organ in which MNV-1 titers were significantly reduced at both 9 and 18 hpi in pIgR KO animals (**see** Figures 1, S1). Host RNA was isolated from the naïve ileum of both groups of mice and analyzed by RT-qPCR for transcript levels of the anti-inflammatory genes TGF-β and IL10, the pro-inflammatory and antiviral genes TNF-α, iNOS, IFN-ɣ, IFN-β, IFN-λ, and ISG15. Both IFN-ɣ and iNOS transcript levels were significantly enhanced in pIgR KO animals compared to WT B6 controls (Figure 4A-B), while transcript levels of inflammatory molecules IFN-β, TNF-α and ISG15 were similar (Figure 4C–E). Transcript levels of TGF-β, a cytokine which promotes IgA class switching (Cerutti 2008), were also similar between strains (Figure 4F). IFN-λ and IL10 were undetectable in either group (data not shown).

**Figure 4:**
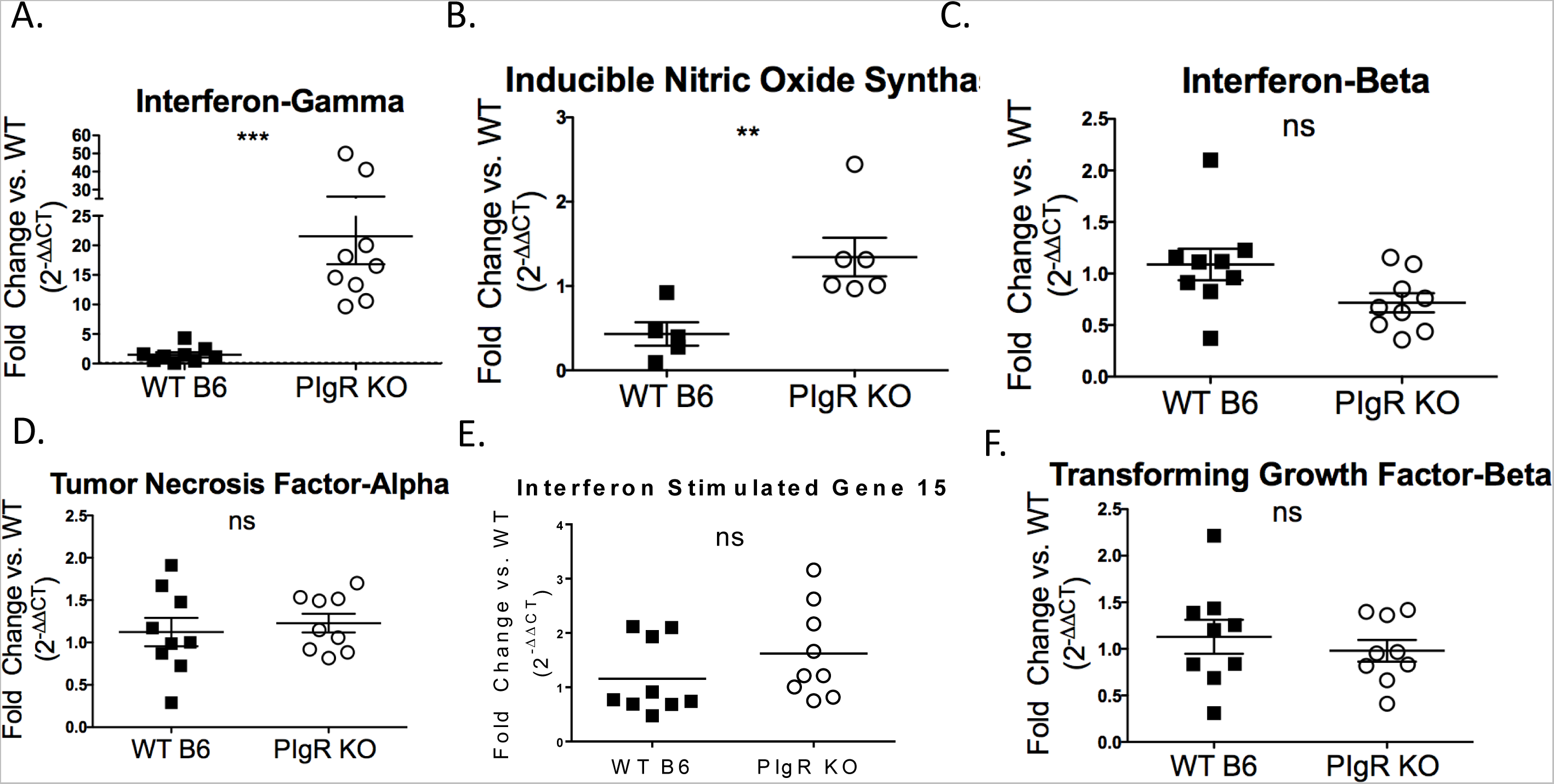
IFN-ɣ and iNOS Levels are enhanced in pIgR KO mice. **A-E.** Total ileal RNA was isolated from naïve pIgR KO and WT B6 mice to determinehost gene levels in reference to *gapdh*. Data is shown in reference to WT B6 and displayed as 2^∧^-(ΔΔCT). ΔCT values were analyzed for significance. Each symbol is the datapoint from an individual animal. The data mean is indicated for each group. *= P<0.05. **= P<0.01 ***= P<0.001. ****= P<0.0001. ns = not statistically significant.

These data demonstrate that pIgR KO animals have enhanced levels of iNOS and IFN-ɣ, two well-known molecules with anti-viral activities (Karupiah, Xie et al. 1993, Persichini, Colasanti et al. 1999, Saura, Zaragoza et al. 1999, Samuel 2001). No significant mortality is seen in iNOS-deficient mice infected with MNV-1 (Karst, Wobus et al. 2003), and MNV-1 replicates similarly in bone marrow-derived macrophages from WT and iNOS-deficient mice (Wobus, Karst et al. 2004). On the other hand, IFN-ɣ is a cytokine known to inhibit MNV-1 translation (Changotra, Jia et al. 2009, Maloney, Thackray et al. 2012). Thus, these findings suggest that elevated levels of some antiviral cytokines, in particular IFN-ɣ, may limit MNV-1 replication in pIgR KO mice.

### IFN-ɣ and iNOS levels are similar in germ-free pIgR KO and WT B6 mice

To test whether enhanced levels of some antiviral molecules limit MNV infection in pIgR KO mice, we first sought to identify a condition, in which the levels of IFN-ɣ and iNOS are similar between pIgR KO and WT mice. Since SIgA dampens the inflammatory signals of commensal bacteria (Corthesy 2007), we hypothesized that germ-free animals might represent one such condition. Thus, both mouse strains were rederived germ-free and transcript levels from the ileum of these mice infected with MNV-1 for 18 hpi were analyzed. As hypothesized, ileal IFN-ɣ and iNOS transcript levels were no longer statistically different without bacterial stimulation (Figure 5A-B). Similar to conventionally housed animals, IFN-β and TNF-α transcript levels were equivalent between germ-free mice (Figure 5C–D). These data demonstrate that removal of the microbiota equalizes small intestinal IFN-ɣ and iNOS levels in pIgR KO and WT B6 mice, highlighting the immunomodulatory functions of SIg.

**Figure 5:**
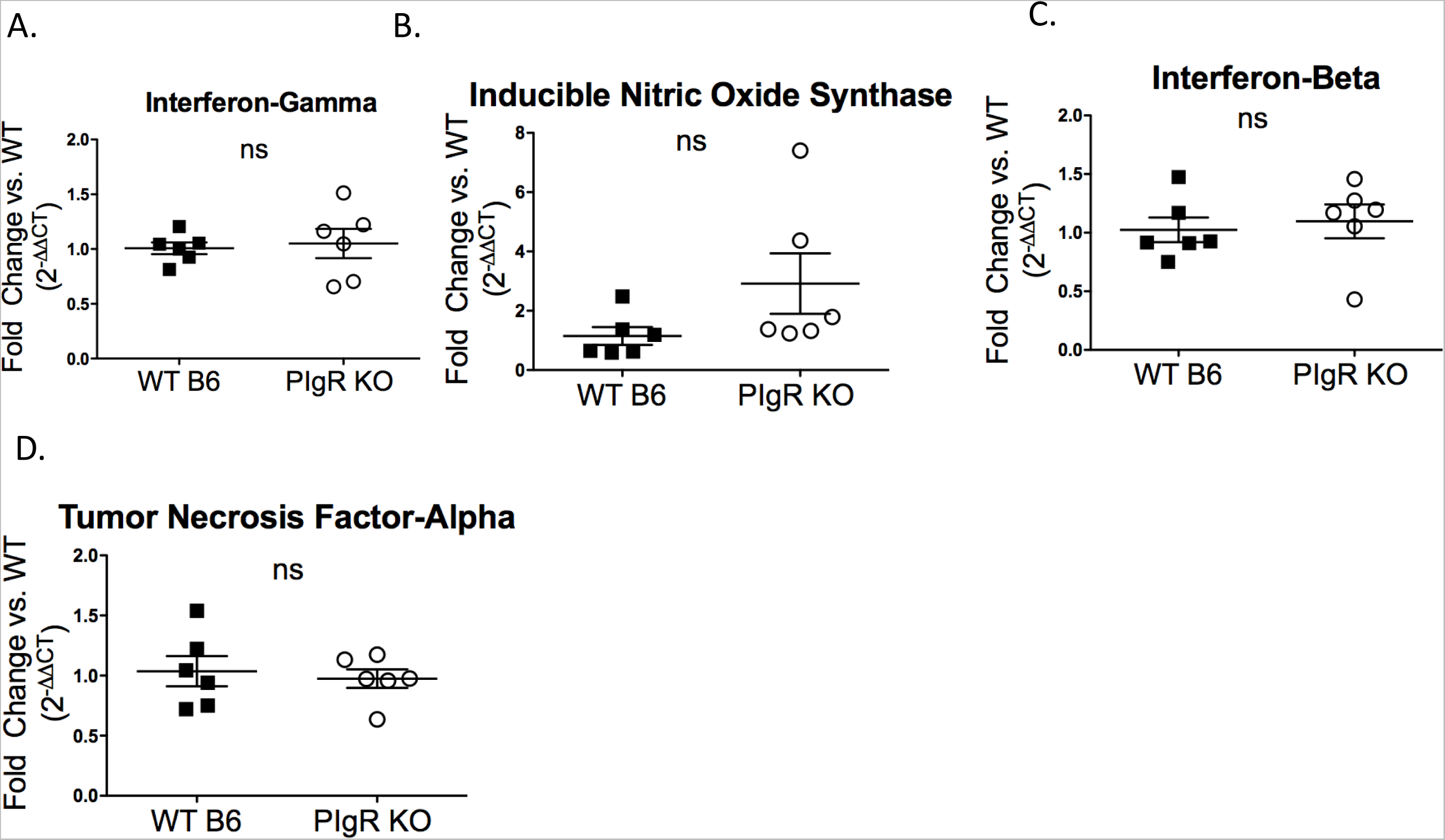
IFN-ɣ and iNOS Levels are reduced in germ-free pIgR KO mice. **A-D.** Total ileal RNA was isolated from germ-free pIgR KO and germ-free WT B6animals infected for 18 hours via o.g. with 3.81x10^5^ PFU/animal MNV-1-NR in 100 μl. Host gene levels were normalized to *gapdh*. Data is shown in reference to WT B6 and displayed as 2∧-(ΔΔCT). ΔCT values were analyzed using unpaired Student’s T test. Data are from two independent experiments; each symbol is the datapoint from an individual animal. The data mean is indicated for each group. ns = not statistically significant.

### MNV-1 infection is similar in germ-free pIgR KO and WT B6 mice

The previous findings suggest that, in the absence of pIgR and hence SIg, inflammatory responses to enteric bacteria may create an inhospitable environment for MNV-1 replication. To test this hypothesis, we infected germ-free pIgR KO and WT animals with MNV-1-NR and assessed replicated viral titers via plaque assay at 18 hpi. Consistent with our hypothesis, no differences were observed in replicated viral titers throughout the entire gastrointestinal tract (i.e., duodenum, jejunum, ileum, cecum, colon) and draining mesenteric lymph nodes in germ-free pIgR KO animals compared to germ-free WT B6 controls (Figure 6). These data demonstrate that MNV-1 replicates similarly in pIgR KO and WT mice under conditions were inflammatory markers such as IFN-ɣ and iNOS are equivalent. These data suggest that pIgR and SIg do not directly inhibit MNV infection but that SIg promote norovirus infection through the maintenance of intestinal homeostasis.

**Figure 6:**
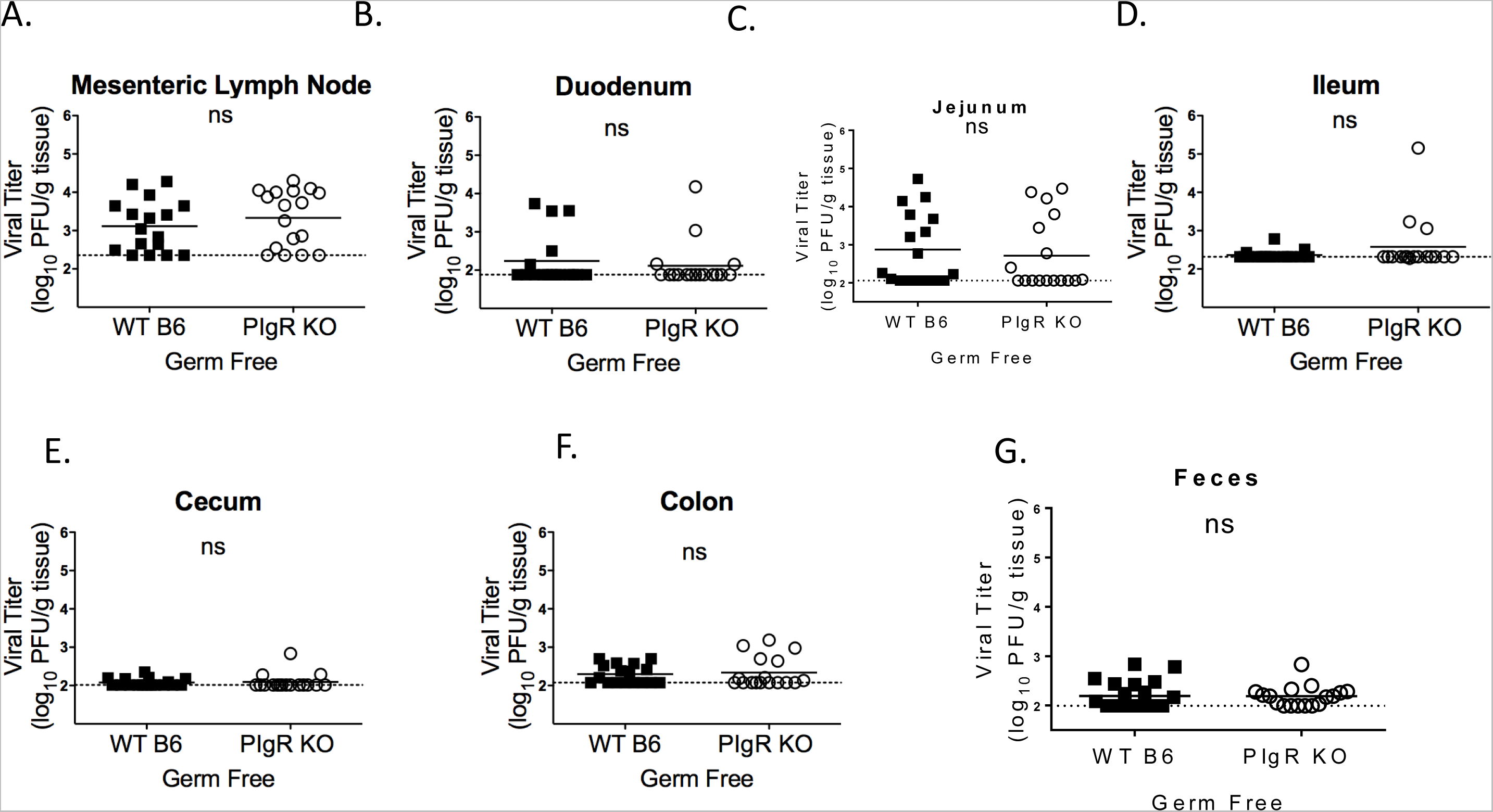
Equalizing IFN-g levels equalizes norovirus infection. **A-G.** Germ-free animals were infected by o.g. with 3.8×10^5^PFU/animal MNV-1-NR in100 μl, and tissues were harvested at 18 hpi in a darkened room using a red photolight. The tissue homogenate was serially diluted and exposed to light for 30 min. Replicated viral titers in the indicated tissue of germ-free WT B6 and pIgR KO animals were assessed via plaque assay. The limit of detection for each graph was as follows (log PFU/g tissue): **A.** 2.356 **B.** 2.881 **C.** 2.057 **D.** 2.317 **E.** 2.014 **F.** 2.077 **G.** 1.993. Data are from four independent experiments and each symbol is a datapoint from an individual animal. Error bars are standard error of the mean (SEM). Values were analyzed using unpaired Student’s T test. ns = not statistically significant.

## Discussion

SIg provides a first line of defense against several enteric pathogens by reducing immunogenicity of lumenal bacterial communities and mediating immune exclusion (reviewed in (Corthesy 2007)). PIgR and SIg are capable of neutralizing intraepithelial pathogens and mediate expulsion from the host (reviewed in (Kaetzel 2005, Corthesy, Benureau et al. 2006). SIg can also bind and neutralize pathogens that have crossed the epithelial barrier and excrete them back into lumen (Corthesy 2007). Surprisingly, despite the ability to bind recombinant, non-MNV specific dIgA, SC and SIgA, removing SIg-mediated innate intestinal defenses in mice did not alter MNV-1 binding to the FAE or internalization into PP. Instead, our study demonstrated that pIgR KO mice, which lack SIg, showed reduced MNV-1 infection. This suggests natural SIg does not exclude MNV from the intestine or mediate intraepithelial expulsion of this virus. More importantly, these data indicate that natural SIg do not protect from norovirus infection. This result is surprising considering that other enteric pathogens are controlled in this manner (Davids, Palm et al. 2006, Schwartz-Cornil, Benureau et al. 2002, Wijburg, Uren et al. 2006). To the best or our knowledge, this represents the first reported example in which pIgR and SIg have proviral functions and promote infection by an enteric pathogen.

Further characterization of pIgR KO mice indicated that reduced infection in pIgR KO mice was not due to a lack of target cells. Instead, these mice had ~2.5 fold greater numbers of macrophages, DCs and B cells in their PP, indicating pIgR KO mice have enlarged PP. In addition, the lack of norovirus infection was also not due to a lack of M cells, suggesting that M cell development is independent of SIg. However, analysis of host mRNA levels in the small intestine revealed that IFN-ɣ, a cytokine that inhibits MNV translation (Changotra, Jia et al. 2009) and iNOS, were elevated in naïve pIgR KO animals compared to WT controls. Enhanced IFN-ɣ and iNOS transcript levels in pIgR KO mice were dependent on the presence of the microbiota. Germ-free pIgR KO and WT B6 mice had equivalent cytokine and viral loads. Therefore, these data suggest that the lack of natural SIg indirectly inhibited MNV-1 infection.

Our data further indicate that the immune response generated towards the enteric microbiota promotes norovirus infection. This is consistent with previous work, which demonstrates that norovirus infections are modulated directly and indirectly by the presence of commensal bacteria (Karst 2016). In case of MNV, infection of antibiotic-treated mice results in decreased MNV loads in the ileum (Jones, Watanabe et al. 2014, Kernbauer, Ding et al. 2014), and commensal bacteria promote MNV persistence in the intestine via modulation of type III interferon responses (Baldridge, Nice et al. 2015). The study described herein represents another example by which enteric bacteria indirectly influence MNV pathogenesis. Specifically, MNV infection was reduced only in conventional pIgR KO mice, which exhibited elevated levels of iNOS and IFN-ɣ, but not in their germ-free counterpart, when cytokine levels were similar to WT mice. Previous studies have also reported enhanced inflammatory activation in pIgR KO mice, resulting in lethal systemic hyperactivity (Karlsson, Johansen et al. 2010), spontaneous COPD (Richmond, Brucker et al. 2016), and enhanced susceptibility to chemically-induced colitis (Murthy, Dubose et al. 2006). A recent study provides a possible explanation based on elevated serum IgA levels in pIgR KO mice (Johansen, Pekna et al. 1999, Shimada, Kawaguchi-Miyashita et al. 1999). Specifically, the study determined that innate immune cell detection of serum IgA via the IgA receptor (FcαR1) in conjunction with pattern recognition receptor activation, resulted in synergistic enhancement of pro-inflammatory cytokine production and release (Hansen, Hoepel et al. 2017). Another study demonstrated greater levels of enteric bacteria in the MLN of pIgR KO mice and highly specific serum IgA responses to several enteric bacterial communities (Sait, Galic et al. 2007). Therefore, enhanced serum IgA levels, coupled with enhanced access of immune cells to bacteria may result in this improperly controlled immune response to enteric commensals, which we and others have observed in pIgR KO mice.

Interestingly, the observed increase in inflammatory immune responses was not uniform, *i.e.*, elevated iNOS and IFN-ɣ levels in conventional pIgR KO mice, but no change in transcript levels of other pro-and anti-inflammatory mediators (see Fig. 4). The reason for this remains unclear. One potential explanation may come from the observed increases in the number of APC subsets (DCs, macrophages and B cells) in the PP of pIgR KO animals compared to WT controls. These cell types have both FcαR1 and pathogen recognition receptors and are capable of pro-inflammatory cytokine secretion (Sakamoto, Shibuya et al. 2001, Otten, Groenveld et al. 2006, Hansen, Hoepel et al. 2017). In particular macrophages are known to secrete iNOS and IFN-ɣ (Munder, Mallo et al. 1998, Cherdantseva, Potapova et al. 2014). Previous studies have also found that intra-epithelial lymphocytes (IELs) are enhanced in the small intestines of pIgR KO mice (Yamazaki, Shimada et al. 2005, Kato-Nagaoka, Shimada et al. 2015). Activated IELs from pIgR KO animals exhibited more abundant IFN-ɣ secretion than WT IELs (Kato-Nagaoka, Shimada et al. 2015), pointing to another potential source for the enhanced IFN-ɣ in these animals. Of note, activation of IELs via the T cell receptor *in vivo* results in significant reduction of MNV viral titers in the small intestine and MLN of WT B6 mice (Swamy, Abeler-Dorner et al. 2015), suggesting a potential cell type that may mediate the antiviral activity in pIgR KO mice.

Collectively, our findings point to a model in which the immunomodulatory functions of SIg aid in norovirus replication. Interaction of noroviruses with SIg complexes could skew the inflammatory viral immune response since uptake of SIgA by mucosal DCs promoting tolerogenic responses and intestinal homeostasis (Corthesy 2007). This might reduce viral clearance and immune memory formation and account for the weak inflammatory response and poor lasting immunity observed in norovirus infections (Newman and Leon 2015). Another potential model of how SIg may promote norovirus infection could come from co-infection of MNV-susceptible APCs when these cells take up several virions via multivalent SIg complexes. Co-infection can increase viral fitness via complementation or recombination (e.g., (Allison, Thompson et al. 1990, Chen, Du et al. 2015, Erickson, Jesudhasan et al. 2017)).

Taken together, our study demonstrates that contrary to previous studies SIg do not protect from MNV infection but that instead the presence of pIgR and SIg promotes infection. Thus, these molecules are host factors that can have either anti- or pro-microbial functions depending on the pathogen. In addition, these findings represent another example in which the microbiota modulates enteric virus pathogenesis. Lastly, these data further highlight that MNV as a natural mouse pathogen has optimized its infection under conditions of intestinal homeostasis. When these conditions are perturbed, such as the absence of pIgR, SIg or enteric bacteria, virus infection is compromised. In the future, it will be interesting to test whether this holds true for other enteric viral infections.

## Acknowledgements

We are indebted to Dr. Thaddeus Stappenbeck for the generous gift of pIgR KO breeders and Dr. Kathryn Eaton and the germ-free mouse core at the University of Michigan. This work was funded in part by NIH R21 AI103961 to C.E. W. and the University of Michigan Host-Microbiome Initiative. H.T. was supported in part by a University of Michigan Rackham Merit fellowship and NIH training grants T32AI007413 and T32DK094775.

## Supplemental Figures

**Supplemental Figure 1: Kinetics of MNV-1-Neutral Red infection in C57BL/6 mice A-H.** WT B6 animals were infected by o.g. with 3.8×10^5^PFU/animal MNV-1-NR in 100 μl and tissues of five mice were harvested at the indicated timepoints in a darkened room using a red photolight. The tissue homogenate was serially diluted, and exposed to light for 30 min. Replicated viral titers in the indicated tissues were assessed via plaque assay. Error bars are standard error of the mean (SEM).

**Supplemental Figure 2: Characterization of pIgR KO animals A-B.** Serum and feces was collected from naïve pIgR KO and WT B6 and IgA concentrations were measured via ELISA. **A.** Serum diluted 1:500. **B.** 10% fecal slurries diluted 1:200. Each symbol is a datapoint from an individual animal. Error bars are standard error of the mean (SEM). Data were analyzed using unpaired Student’s T test. *= P<0.05. **= P<0.01 ***= P<0.001. ****= P<0.0001. nd = not detected.

**Supplemental Figure 3: Acute norovirus infection is reduced in pIgR KO mice A-G.** Animals were infected o.g. with 3.81x10^5^PFU/animal MNV-1-NR, and tissues were harvested at 9 hpi in a darkened room using a red photolight. The tissue homogenate was serially diluted and exposed to light for 30 min. Replicated viral titers in the indicated tissue of WT B6 and pIgR KO animals were assessed via plaque assay. The limit of detection for each graph was as follows (log PFU/g tissue): **A.** 2.104 **B.** 2.26 **C.** 2.338 **D.** 2.525 **E.** 2.313 **F.** 2.216 **G.** 2.507. Data are from three independent experiments and each symbol is a datapoint from an individual animal. Error bars are standard error of the mean (SEM). Data were analyzed using unpaired Student’s T test. *= P<0.05. **= P<0.01 ns = not statistically significant.

**Supplemental Figure 4: Natural secretory immunoglobulins do not aid in MNV access to the Peyer’s patch A.** ELISA was performed by coating microtiter plates with 0.1 mg/ml of recombinant murine secretory IgA (SIgA), murine polymeric IgA (pIgA), or murine secretory component (SC) and incubated with bacterially expressed protruding- (P) or shell- (S) domains of the MNV-1 capsid or ELISA coating buffer (B). Protein domains were detected using an MNV-1 capsid antibody followed by a peroxidase-conjugated secondary antibody. Absorbance was read at 405 nm. Six replicates are shown. Error bars represent standard error of the mean (SEM). **B.** Intestinal ileal loop assay in WT B6 and pIgR KO. 100μl (3.81×10^5^PFU/loop) MNV-1was injected into the closed loop and incubated for 25 min. Viral genome copy equivalents were measured by RT-qPCR in Peyer’s patches. Data are from four independent experiments and each symbol is a datapoint from an individual PP. The data mean is indicated for each group. Data were analyzed using unpaired Student’s T test. ns = not statistically significant.

